## Supplementary Figures for "Antibiotic persistence of *Brucella abortus* in its protective intracellular niche"

### Supplementary figures and legends:

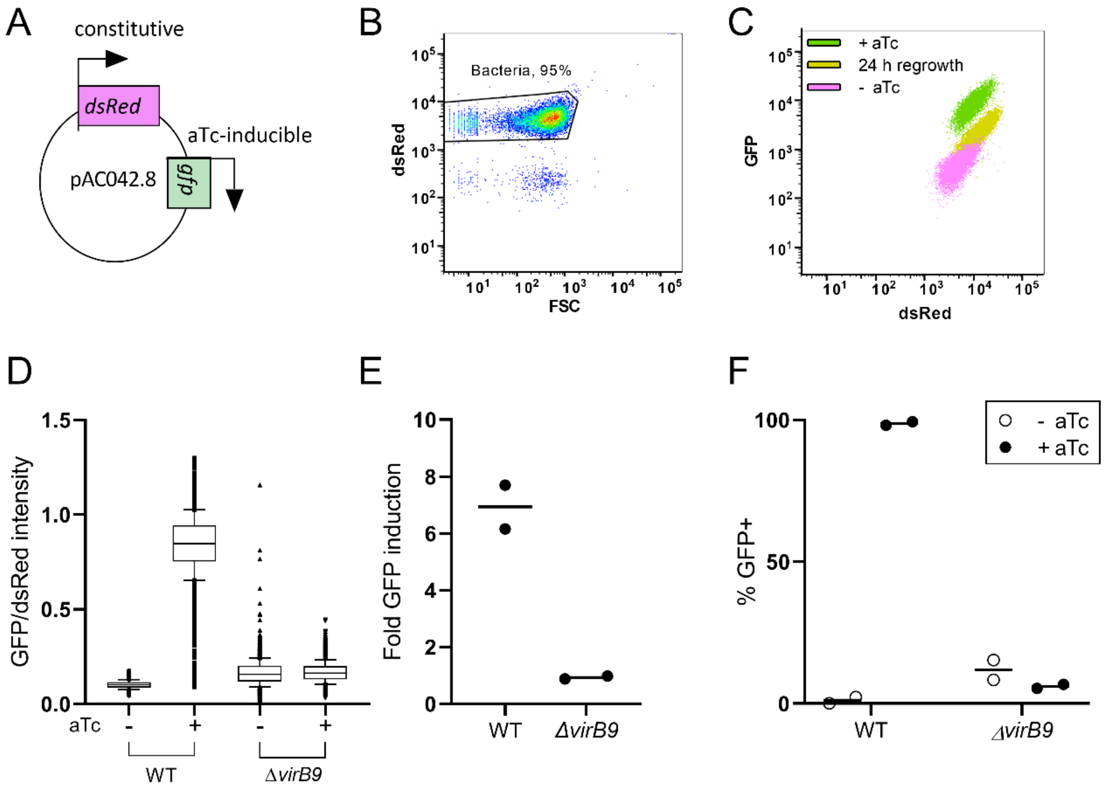


**Figure S1: Response of *B. abortus* pAC042.8 to inducer in broth and in RAW macrophage infection.** (A) Schematic representation of plasmid pAC042.8. Bacteria carrying pAC042.8 constitutively express dsRed and express GFP under a tetracycline-inducible system. (B) Flow-cytometric identification of the bacterial population grown till the exponential phase based on its dsRed fluorescence and forward-scatter (FSC) property. (C) Flow-cytometric detection of the GFP fluorescence in the bacterial population grown till exponential phase (magenta), after the addition of the inducer (green) and 24 h after removal of the inducer (yellow). For (B) and (C), displayed is a representative dot blot of one experiment of at least 3 independent replicates. (D, E, F) RAW macrophages were infected with *B. abortus* pAC042.8 or *B. abortus* Δ*virB9* pAC042.8 for 27 h. Cells were treated with aTc (100 ng/ml) at 23 hpi for 4h before fixation and imaging. (D) Box plot showing the size-independent response of the bacterial population to the inducer. (E) Fold induction of the bacterial population response to the inducer. Fold induction was calculated by dividing the median values of the induced population by the non-induced one between matching conditions. Each dot represents one independent replicate, the horizontal line represents the mean, n=2. (F) Percentage of GFP+ infection sites. Horizontal bars represent the mean, n=2.

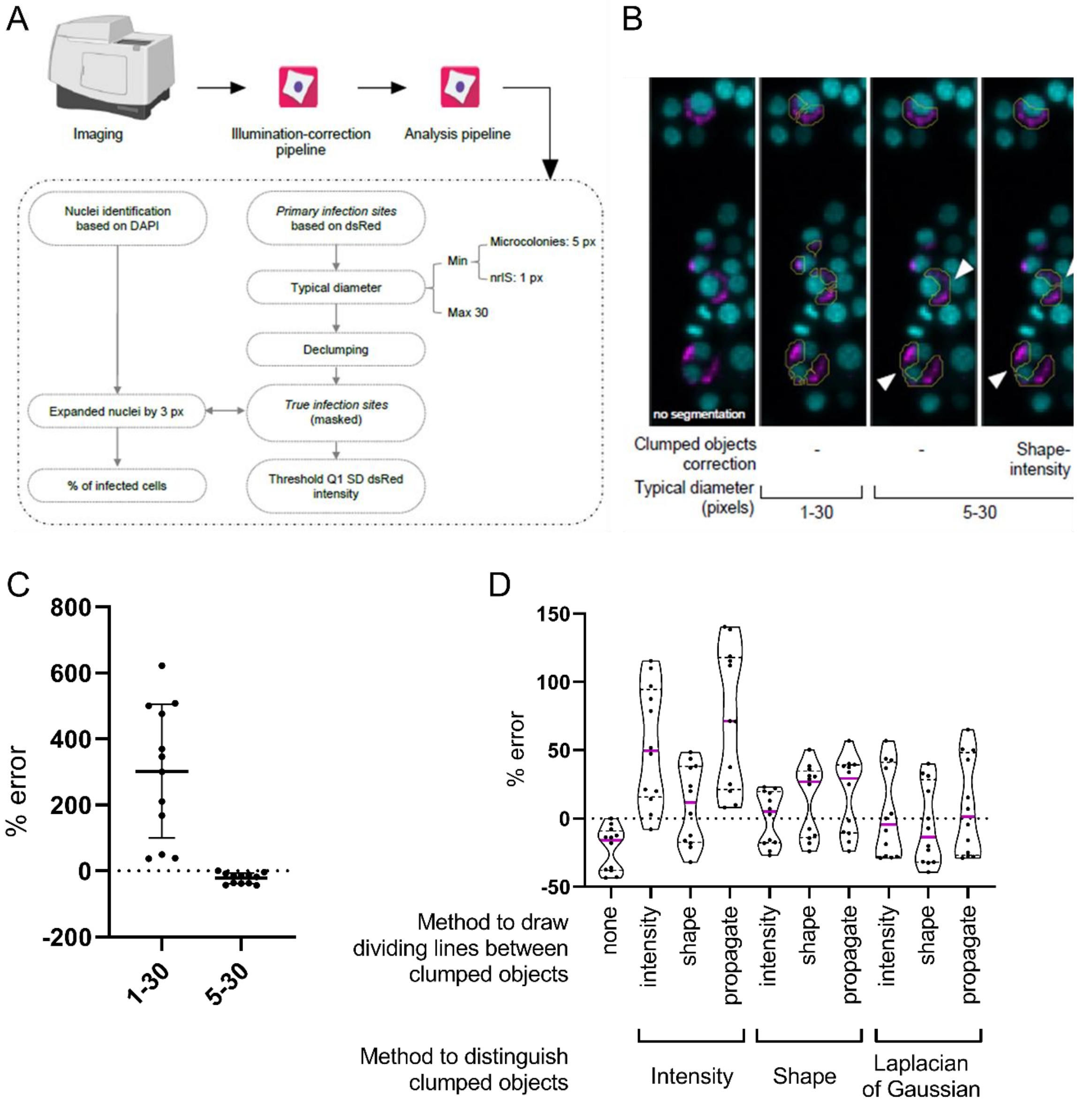


**Figure S2: Visual overview of CellProfiler workflow and method to distinguish clumped objects.** (A) Image analysis workflow for CellProfiler for the segmentation of nuclei and infection sites. The minimum typical diameter was set to different values in the pipelines to either identify non-replicative infection sites or microcolonies. Nuclei were expanded by 3 pixels and only infection sites residing inside the expanded nuclei were kept for the analysis. (B) Representative images from RAW264.7 macrophages infected for 27 h with *B. abortus* pAC042.8 constitutively expressing dsRed (magenta). Macrophage nuclei were stained with DAPI (cyan). Image analysis was performed with CellProfiler to segment nuclei and bacteria and to extract measurements. The left panel shows microcolonies segmentation with a typical diameter set between 1 and 30 pixels. The middle and right panels show microcolonies segmentation with a typical diameter between 5 and 30 pixels, without (middle) clumped-objects correction. The right panel shows microcolonies segmentation after identification of clumped objects based on the shape followed by the division of those based on the intensity. Arrows indicate examples of separation of merged microcolonies. (C) Percent error between manual count and CellProfiler count using a typical diameter between 1 and 30 or between 5 and 30. Data represent the mean ±SD, n=10 (5 x 2 sites per experiment). (D) Violin plot showing the distribution of the percent error between manual count and CellProfiler count of microcolonies using several methods to distinguish and separate clumped objects. The pink line represents the median, dotted lines represent quartiles. n=10 (5 x 2 sites per experiment).

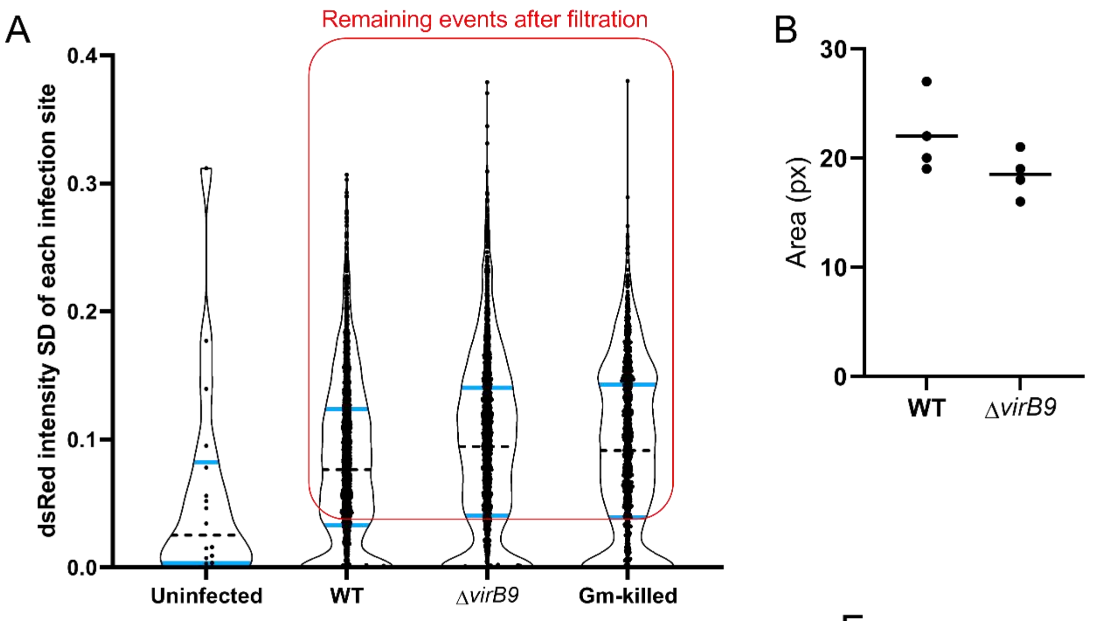


C

| P90 | WT | ΔvirB9 |
|---|---|---|
| Area (px) | 22.00 | 18.50 |
| Min-max | 19.5-24.5 | 17-20 |
| Typical diameter (px) | 5.29 | 4.85 |
| Min-Max | 4.98-5.59 | 4.65-5.05 |

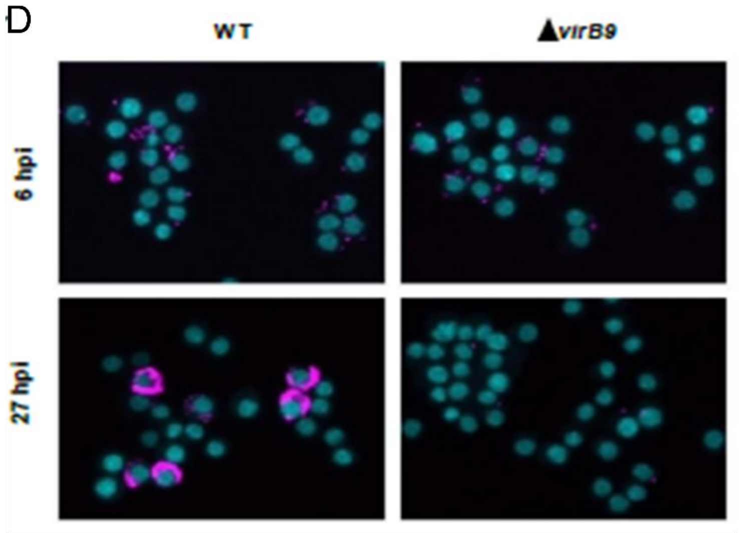


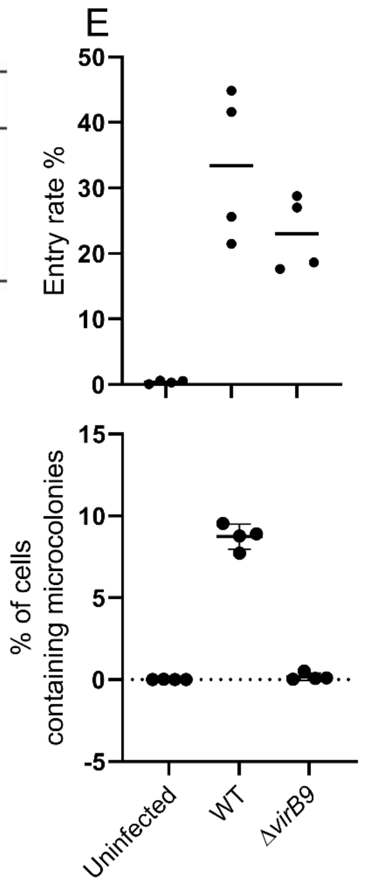


**Figure S3: CellProfiler pipeline set-up to define infection site sizes.** (A) Representative violin plot depicting the distribution of the standard deviation of the dsRed intensity for each infection site of *B. abortus* wild-type (WT), Δ*virB9* and wild-type killed with gentamicin for 24 h (Gm-killed) all carrying pAC042.8. Horizontal blue lines display first and third quartiles, horizontal black dotted line represents median. Red box shows remaining event after filtration with CellProfiler pipeline. n=2 (B) Area of infection sites before intracellular replication. The $90^{th}$ percentile (P90) of the measured area for each condition after 5 h infection. Horizontal lines represent the average, n=4 (2 x 2 technical replicates). (C) Average area and typical diameter of infection sites of *B. abortus* wild-type (WT) and Δ*virB9*. Table shows the equivalence between the measured area and the diameter of a circle using the formula $dd=2(\sqrt{A}/\sqrt{\pi})$. Displayed are the averages from 2 technical replicates from 2 independent experiments. (D) Representative images from RAW264.7 infected for 6 h or 27 h with *B. abortus* WT or Δ*virB9*. Bacteria constitutively expressed dsRed (magenta) and macrophage nuclei were stained with DAPI (cyan). (E) Entry rate (top) and percentage of cells containing microcolonies (bottom) from macrophages infected with *B. abortus* WT or *B. abortus* Δ*virB9* carrying plasmid pAC042.8. The entry rate and the percentage of cells containing microcolonies were determined using the number of nuclei associated with at least one non-replicative infection site or one microcolony in the 3 pixels-expanded nuclei area. Horizontal lines show the average, n=2 (2 x 2 technical replicates).

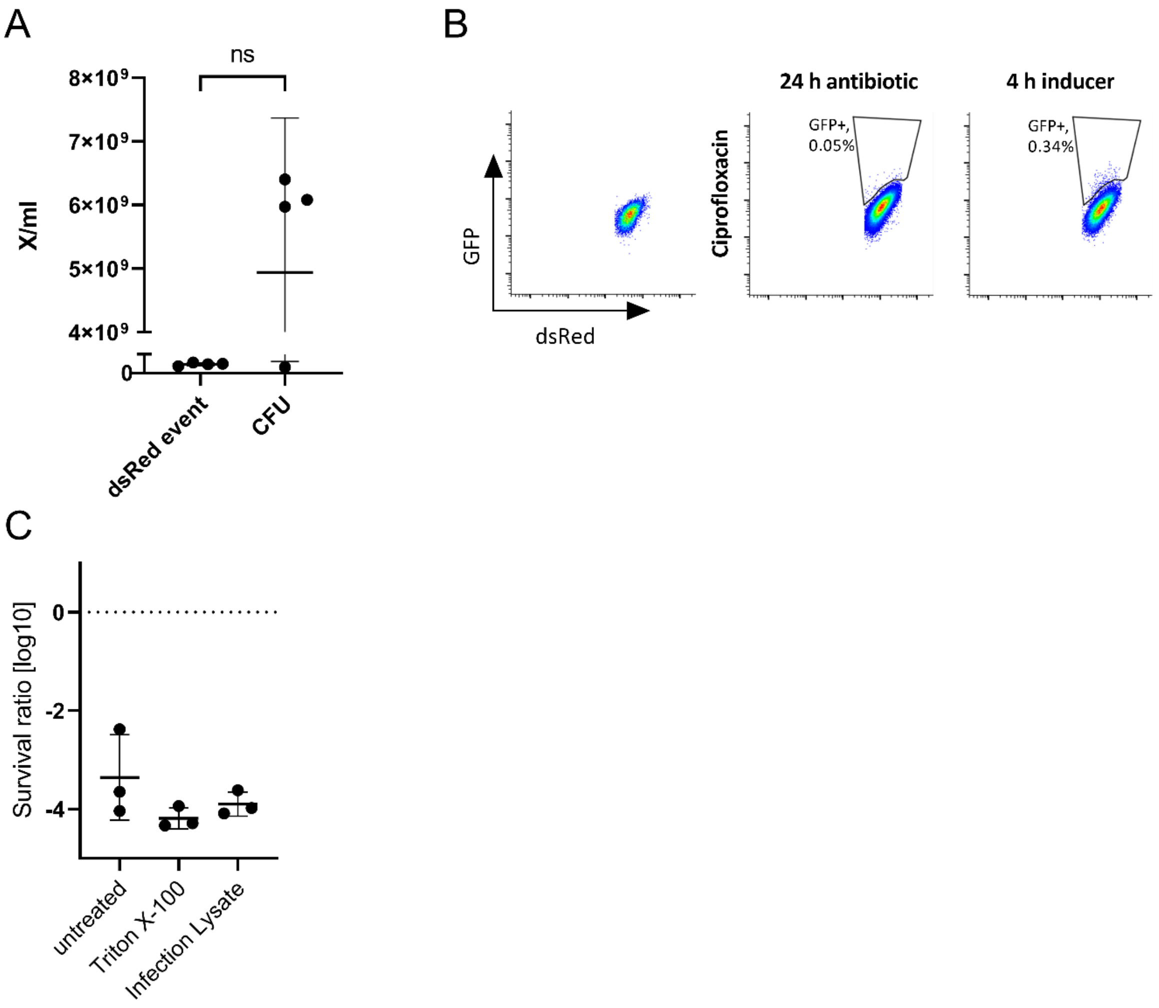


**Figure S4: GFP-induction in broth, control treatments for macrophage lysis, and pre-dilution of inoculum.** (A) Comparisons between CFU counts and flow cytometric counts using counting beads. Data represent the mean ± SD, n=4. Statistical analysis was performed using paired t-test (ns, non-significant $p > 0.05$). (B) Flow-cytometric identification of the GFP+ population. Representative dot blots of one experiment of 3 independent replicates. (C) *B. abortus* was grown to mid exponential phase, macrophages were infected with *B. abortus* for 10 min, and *B. abortus* was incubated for 10 min in Triton X-100. After each step samples were washed, and one part of the sample was used to enumerate CFU/ml before ciprofloxacin treatment. Another part of each sample was subcultured in TSB plus ciprofloxacin and after 24 h CFU/ml were enumerated. The survival ratio was calculated by division of CFUs recovered before by CFUs recovered after ciprofloxacin treatment.
